# Discovery of Novel Bilaterian Signaling Peptides Using Cone Snail Toxins

**DOI:** 10.1101/2022.08.05.502922

**Authors:** Thomas Lund Koch, Joshua P. Torres, Robert P. Baskin, Paula Flórez Salcedo, Kevin Chase, Baldomero M. Olivera, Helena Safavi-Hemami

**Affiliations:** Department of Biomedical Sciences, University of Copenhagen, Copenhagen-N, 2200, Denmark; School of Biological Sciences, University of Utah, Salt Lake City, UT 84112, USA; The Ohio State University College of Medicine, Columbus, OH, 43210, USA; Department of Neurobiology, University of Utah, Salt Lake City, UT 84112, USA; Department of Biochemistry, University of Utah, Salt Lake City, UT 84112, USA

## Abstract

Peptide hormones and neuropeptides form a diverse class of signaling molecules that control essential processes in animals. Despite several breakthroughs in peptide discovery, many signaling peptides remain undiscovered. Recently, we demonstrated the use of somatostatin-like toxins from cone snail venom to identify homologous signaling peptides in prey. Here, we demonstrate that this toxin-based approach can be systematically applied to the discovery of other unknown bilaterian signaling peptides. Using large sequencing datasets, we searched for homologies between cone snail toxins and putative peptides from several important model organisms representing the snails’ prey. We identified five toxin families that share strong similarities with previously unknown signaling peptides from mollusks and annelids. One of the peptides was also identified in rotifers, brachiopods, platyhelminths, and arthropods, and another was found to be structurally related to crustacean hyperglycemic hormone, a peptide not previously known to exist in Spiralia. Based on several lines of evidence we propose that these signaling peptides not only exist but serve important physiological functions. Finally, we propose that the discovery pipeline developed here can be more broadly applied to other systems in which one organism has evolved molecules to manipulate the physiology of another.

## Introduction

Neuropeptides and peptide hormones (collectively referred to as signaling peptides) are important signaling molecules found throughout the Animal Kingdom (Jékely 2013; Koch and Grimmelikhuijzen 2019; Nikitin 2015). Signaling peptides are expressed in multiple tissues and cell types, such as neurons, endocrine glands, and other secretory cells. Evolutionarily, many signaling peptides are ancient, and their origin can sometimes be traced back to the common ancestor of protostomes and deuterostomes, the two major groups of bilaterians (Jékely 2013; Mirabeau and Joly 2013). Examples include oxytocin/vasopressin-like peptides, the insulin family, and neuropeptide Y peptides.

Most signaling peptides are 5-50 amino acids in length and released from larger precursor molecules by the action of specific proteases that typically cleave at basic or dibasic amino acid residues. The precursors contain an N-terminal signal sequence that targets the peptide to the secretory pathway and typically also contain spacer regions of mostly unknown functions (also referred to as pro-/post-peptide or leader regions). This complex precursor structure of signaling peptides is typically accompanied by contrasting patterns of evolution. The N-terminal signal sequence often diverges substantially, whereas the signaling peptide region and flanking proteolytic processing sites are generally conserved within a single phylum with only minor variations. The spacer regions are typically under genetic drift. Thus, comparative sequence analysis of signaling peptide precursors tends to show a pattern of relaxed purifying, close to neutral selection in the spacer regions and signal sequence, and strong purifying selection in the region encoding the mature signaling peptide (Foster et al. 2019; Koch et al. 2022; Li et al. 2009; Toporik et al. 2014; Williams et al. 2000).

Most known signaling peptides convey their actions through G protein-coupled receptors (GPCRs) (Mirabeau and Joly 2013; Vaudry and Seong 2014). Notable exceptions are insulins, insulin-like growth factors (IGFs), and epidermal growth factors (EGFs) that signal through receptor tyrosine kinases (Lemmon and Schlessinger 2010), and several FMRF-amide-related neuropeptides that interact with a class of degenerin/epithelial sodium channels (DEG/ENaC) (Furukawa et al. 2006; Gründer and Assmann 2015).

Given their importance in animal biology, extensive research programs have attempted to discover and describe signaling peptides and their receptors. Currently, several hundred different signaling peptides are recognized in humans (Foster et al. 2019; Secher et al. 2016; Tai et al. 2020). Still, the endogenous ligands for almost 100 human GPCRs remain unknown (Laschet et al. 2018).

D*e novo* discovery of signaling peptides is difficult. Many of the yet-unknown signaling peptides may not be highly expressed, may be unstable, or may only be expressed in specific cell types or developmental stages. While some signaling peptides have been discovered bioinformatically, this process involves several conceptual challenges such as identifying atypical cleavage sites and distinguishing true signaling peptides from falsely predicted ones. Thus, additional approaches for signaling peptide discovery are needed.

We and others have previously shown that some venomous animals have evolved toxins that specifically mimic the signaling peptides of their prey or predators (Cruz et al. 1987; Eagles et al. 2022; Kanda et al. 2007; Sachkova et al. 2020; Safavi-Hemami et al. 2016). We refer to these peptides as “doppelganger toxins”. Some doppelganger toxins are known to have originated from an endogenous signaling peptide gene that, following recruitment into the venom gland, experienced positive selection to ultimately mimic the related peptide of the prey or predator species (Koch et al. 2022; Sachkova et al. 2020; Safavi-Hemami et al. 2016). This process is accompanied by the generation of novel, advantageous features of the toxin compared to the endogenous peptide it evolved from, such as enhanced stability, receptor subtype selectivity, or faster action (Ramiro et al. 2022; Safavi-Hemami et al. 2016; Xiong et al. 2020).

Since doppelganger toxins typically share some sequence similarity with the signaling peptide they mimic, it is possible to identify these toxins through sequence homology searches. This has for instance led to the discovery of con-insulins; weaponized insulins derived from the endogenous insulin hormone in cone snails that mimic the insulin expressed in fish prey (Safavi-Hemami et al. 2016), and the arachnid toxin Ta1a, which has common ancestry with crustacean hyperglycemic hormone (Undheim et al. 2015). While this homology approach has identified toxins that share sequence similarity to known signaling peptides, it can also, in principle, be “reversed” *i*.*e*., to search for yet-unknown peptides that share homology with toxins.

Anecdotal evidence has shown that this is indeed possible. For example, Bombesin, a peptide from the poisonous secretions of the European fire-bellied frog (*Bombina Bombina*) that stimulates the release of gastrin inspired the search and ultimately the discovery of homologous peptides in mammals (gastrin-releasing peptide (GRP) and neuromedins) (McDonald et al. 1979; Minamino et al. 1983). Similarly, the sea anemone toxin ShK-like1 led to the discovery of the previously unknown signaling peptide Shk-like2 in the nervous system of cnidarians (Sachkova et al. 2020). Additionally, we recently showed that somatostatin-like toxins from cone snails elucidated the presence of a somatostatin signaling system in protostomes (Koch et al. 2022). Here, we hypothesized that this anecdotally reported, toxin-based approach can be used to systematically unravel the existence of unrecognized signaling peptides.

Cone snails and their toxins represent an ideal system for testing the broader feasibility of this approach. *Conus* is a diverse lineage of ∼ 1,000 species of venomous marine gastropods with a large repertoire of hyper-diverse peptide toxins, called conotoxins. Additionally, cone snails have diverse and typically well-described diets ranging from fish to mollusks and annelid worms (Duda et al. 2001; Olivera et al. 2015; Puillandre et al. 2014). This provides a large library of toxins that evolved to specifically target animals belonging to different phyla. Conotoxins were traditionally identified from dissected or milked venom, but the advent of next-generation sequencing has enabled the identification of putative conotoxin sequences from venom gland transcriptome and exome data (Abalde et al. 2020; Li et al. 2018; Phuong et al. 2019; Phuong et al. 2016), revolutionizing the pace of conotoxin discovery. At the same time, comprehensive transcriptomics and genomics datasets of several model organisms representing *Conus* prey have become available, making it possible to systematically test the toxin-based approach proposed here.

By performing a systematic homology search of conotoxins and predicted secreted proteins from cone snail prey, we discover five doppelganger toxin families that share strong similarities with proteins of unknown identity in prey organisms. Based on several lines of evidence, including tissue-specific expression, characteristic evolutionary trace, and structural alignments, we propose that these proteins encode unrecognized bilaterian signaling peptides. Our discovery of these signaling peptides (and their doppelganger toxins) will likely be of importance to neurobiological research as these peptides are found in important model organisms where they are likely to serve critical functions. Finally, our findings serve as a proof of concept for the systematic use of doppelganger toxins for the discovery of yet-unknown signaling peptides. We propose that this approach can be broadly applied to other systems in which one organism has evolved compounds to manipulate the physiology of another. This includes venomous animals and their prey, venomous and poisonous organisms and their predators, and pathogens and parasites and their hosts.

## Results

### Homology searches identify five putative signaling peptides in mollusks and annelids

We investigated the presence of unknown signaling peptides in model organisms from the three phyla of cone snail prey: the chordate *Danio rerio*, the mollusk *Aplysia californica*, and the two annelids *Capitella teleta* and *Platynereis dumerilii* (two annelids were included to have both genomic and transcriptomic data from this phylum) (**Figure 1A)**. Specifically, we aimed to identify unknown signaling peptides that share sequence similarities with cone snail doppelganger toxins (**Figure 1B**). To this end, using published genome and transcriptome datasets we constructed libraries of putative secreted proteins from prey species, which, in principle, should contain all known and unknown signaling peptides (**Figure 1C**). This provided us with a set of 9,328 unique sequences from *D. rerio*, 7,009 sequences from *A. californica*, and 10,659 sequences from the two annelids *C. teleta* and *P. dumerilii* (**Figure 1D**). In addition, we built a database of putative secreted proteins from 92 venom gland transcriptomes of 45 cone snail species. These cone snails belong to 19 phylogenetically diverse clades with different prey preferences (Phylogenetic tree shown in **Supplementary Figure 1**), resulting in a library of 25,989 sequences of conotoxins and conotoxin candidates. This library should principally contain all known and unknown conotoxins.

**Figure 1:**
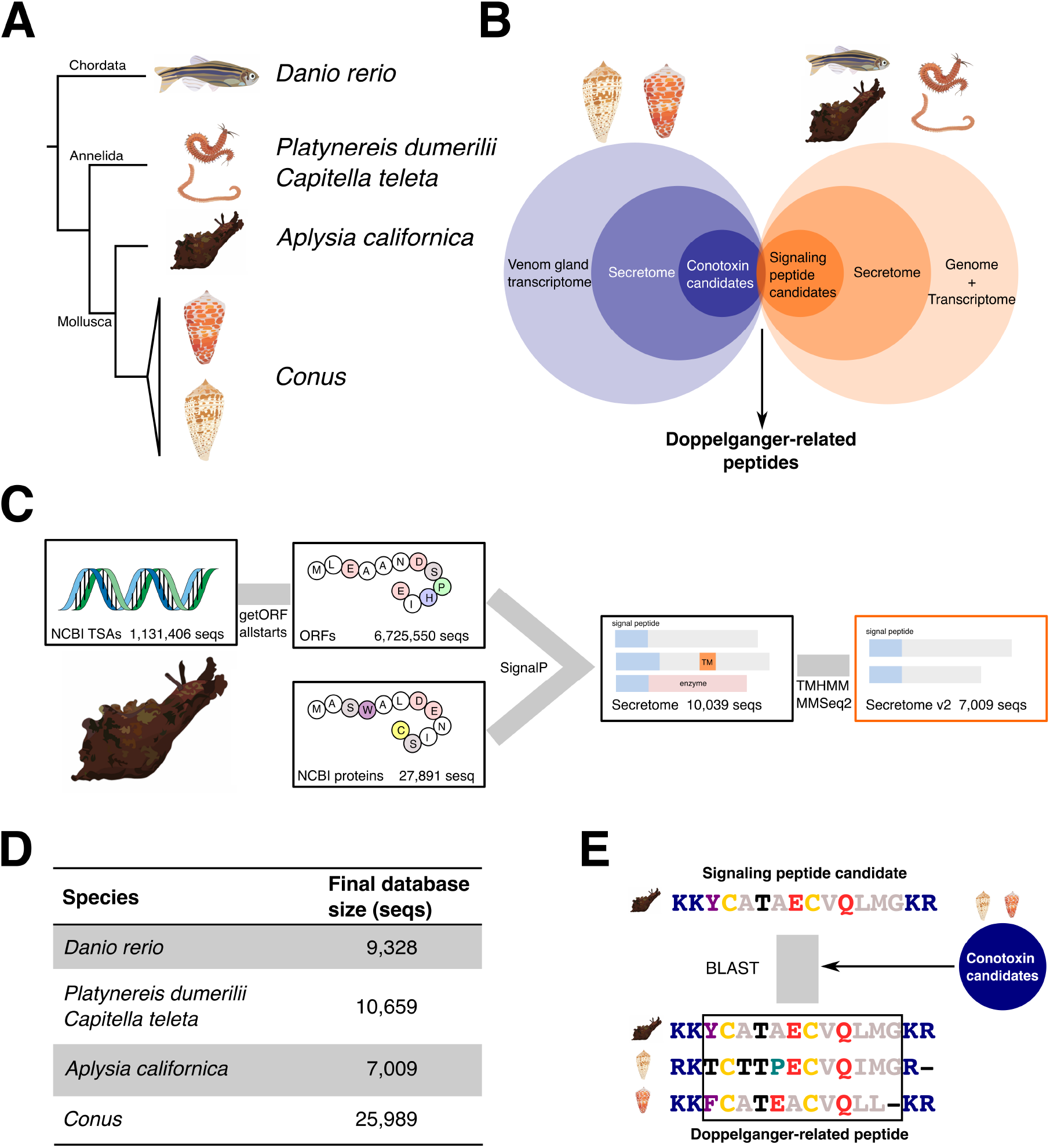
Methodology of toxin-based approach to signaling peptide discovery. (**A**) Model species used in this study representing cone snail prey: *Danio rerio* (zebrafish), the two annelids *Platynereis dumerilii* and *Capitella teleta*, and the mollusk *Aplysia californica* (Californian sea hare) (**B**) The toxin-based approach to signaling peptide discovery is based on finding signaling peptide candidates that share homology with putative conotoxins. (**C**) Workflow for building databases of putative signaling peptide candidates from prey species: predicted proteins and transcriptome assemblies were downloaded and filtered using SignalP. Sequences containing transmembrane domains and those with similarity to known enzymes were removed. A similar approach was used to extract conotoxin candidates from 45 different species of cone snails. (**D**) Resulting database size: 9,328 sequences for *D. rerio*, 7,009 sequences for *A. californica*, and 10,659 sequences for *C. teleta* and *P. dumerilii*. (**E**) Putative prey signaling peptides were blasted against the conotoxin library and retained based on criteria listed under Materials and Methods.

Next, we performed BLAST-based homology searches with the prey databases as queries against the cone snail toxin database (**Figure 1E)**. We employed the following criteria to identify putative new signaling peptides: (1) The protein must be secreted. (2) The prey protein must yield at least two homology hits (e-value > 0.1) to the cone snail toxin database. (3) The prey protein must either have classical signaling peptide processing sites or the peptide must span the entire precursor except for the signal sequence. (4) There must be orthologs in closely related organisms that also show the characteristics of signaling peptide precursors. As we were specifically searching for unknown signaling peptides, we also employed a final criterium: (5) Neither the prey protein nor its orthologs should already have a functional annotation. Using these criteria, we identified five families of sequences from mollusks and annelids that, as we propose, encode novel signaling peptides (**Figure 2**). We refer to these as doppelganger-related peptides (DREPs). DREPs were named based on their sequence or structural characteristics.

**Figure 2:**
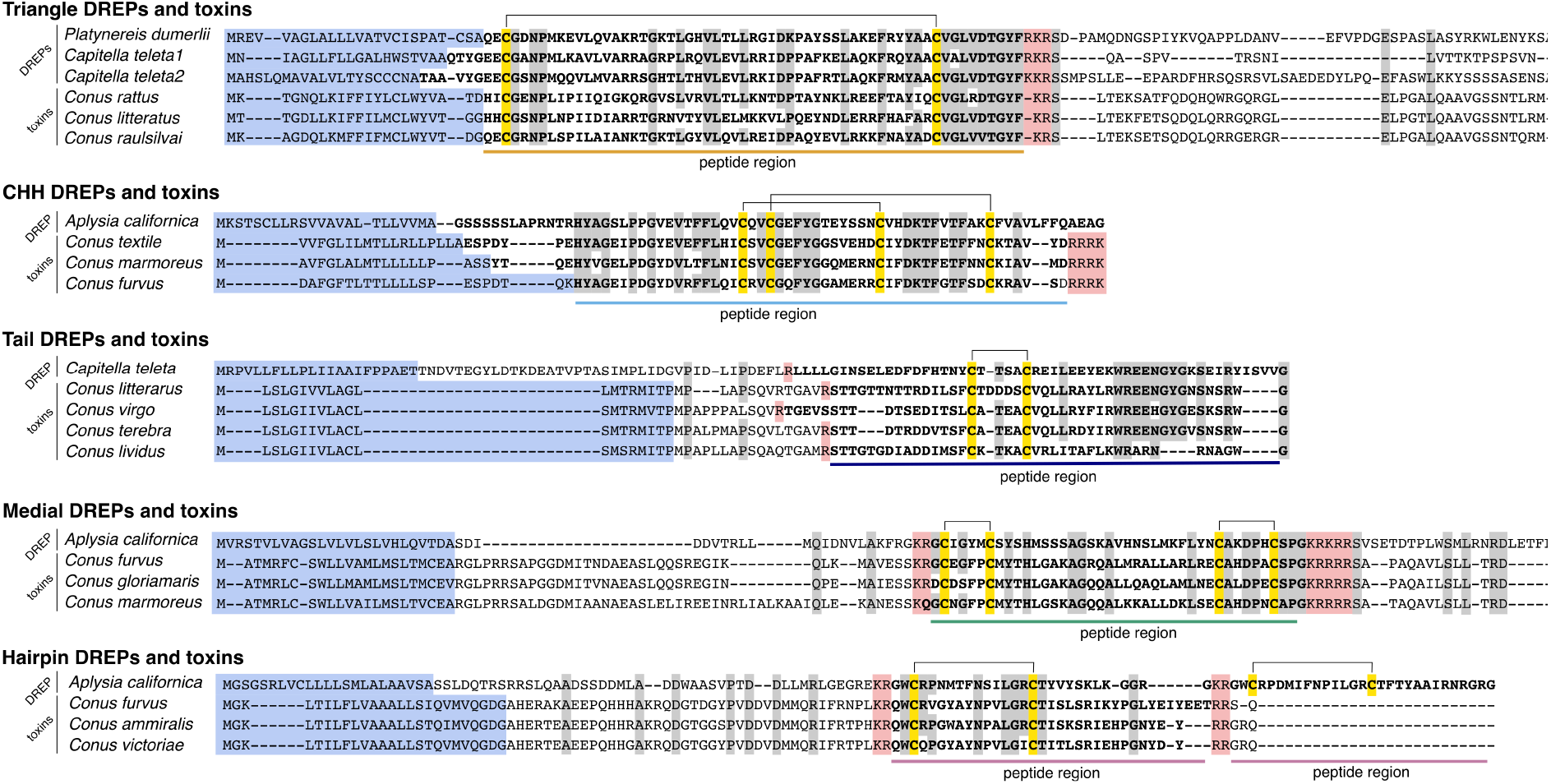
Multiple sequence alignments showing high similarity of the five identified DREP families with doppelganger toxin precursors. Signal sequences are highlighted in blue, cysteines are in yellow with disulfide bonds shown as connecting lines, and processing sites are highlighted in red. Mature DREP and toxin regions are in bold and underlined. Identical amino acids are highlighted in gray.

### Triangle DREPs

The first family of putative signaling peptides, named triangle DREPs, were discovered from a *P. dumerilii* transcript (GenBank ID: HAMN01029001) and two sequences annotated as predicted proteins from *C. teleta* (ELT98797, ELT98795). These showed significant sequence similarity to multiple sequences from the cone snail toxin database (**Figure 2)**. The predicted DREPs and toxins are 59-65 amino acids long and contain a single disulfide bond formed by two cysteines. We queried the toxin database with the initial toxin hits and identified 29 related toxin gene sequences from 11 different cone snail species belonging to the Elisaconus, Africonus, and Rhizoconus clades, all worm-hunting clades (**Supplementary figure 1, Supplementary file 1**). These clades do not form a monophyletic group suggesting that the toxin genes are not expressed or lost in other clades or that the toxins were independently recruited multiple times.

### CHH DREPs

The second DREP family was discovered from toxin hits to a transcript from *A. californica* (GBCZ01041960) (**Figure 2)**. The predicted peptide is 71 amino acids in length, contains two disulfide bonds, and is located immediately downstream of the signal sequence. We identified homologous toxin sequences from the venom gland transcriptomes of the snail hunters *Conus textile, Conus marmoreus*, and *Conus furvus*, belonging to the Cylinder, Conus, and Darioconus clades, respectively (**Supplementary figure 1, Supplementary file 2**). Snail-hunting behavior is believed to have evolved once (Puillandre et al. 2014), and the fact that we only find these toxins in closely related, mollusk-hunting clades suggests that the venom recruitment event happened once in the common ancestor of snail hunters.

### Tail DREPs

The third DREP family was discovered from multiple toxin hits to a sequence annotated as predicted protein from *C. teleta* (ELT87057) (**Figure 2)**. The DREP peptide is 53 amino acids long and is located in the C-terminal region of the precursor. A single disulfide bond is at the N-terminal region of the predicted mature peptide. We identified a total of eight conotoxins with sequence similarity from species belonging to the vermivorous clades of Virgiconus, Africonus, Elisaconus, and Lividoconus (**Supplementary figure 1, Supplementary file 3**).The region located N-terminally to the cysteine loop is enriched in threonine and serine, suggesting that these residues might be post-translationally modified to carry glycosylation as known for other cone snail toxins (Gerwig et al. 2013).

### Medial DREPs

This family was identified based on sequence similarity between multiple toxins to a sequence annotated as predicted protein from *A. californica* (XP_005095677) (**Figure 2)**. The predicted peptide is located in the medial region of the precursor, 40 amino acids in length, and predicted to contain a C-terminal amidation and two disulfide bonds. We note that it is possible that the DREPs encode two peptides rather than a single peptide spanning the entire region. However, there are no conserved processing sites for this cleavage, and we propose processing to a single peptide. We identified 13 toxin sequences in the venom gland transcriptomes that share high similarity to the initial hit from Aplysia. All sequences are from snail-hunting species of the Conus, Darioconus, and Cylinder clades (**Supplementary figure 1, Supplementary file 4**). This suggests that this toxin family evolved in early mollusk-hunters and that these toxins specifically target a molluscan receptor.

### Hairpin DREPs

The final DREP family was identified from hits to an uncharacterized, predicted protein from *A. californica* (XP_005089801) (**Figure 2**). This protein has previously been suggested to encode a signaling peptide based on similarity to a toxin derived from the cone snail *Conus victoriae* from the Cylinder clade, contryphan-Vc1 (Robinson et al. 2016). Interestingly, the Aplysia precursor possesses two copies of similar peptides on the same precursor. The presence of numerous highly similar copies of a single peptide is a common feature of many protostome signaling peptide precursors. Here, we identified similar toxins in *C. furvus* and *C. ammiralis*, two snail-hunting species from the Darioconus and Cylinder clade, respectively (**Supplementary figure 1, Supplementary file 5**). As previously observed, the signal sequence of these toxins is similar to that of the contryphans/O2 superfamily of conotoxins (Robinson et al. 2014). However, apart from the signal sequence and the presence of a single disulfide bond the doppelganger toxin family shares little similarity with contryphans (Contryphans have a much shorter disulfide loop and the mature contryphan toxin is located at the N-terminus of the toxin precursor). Consistent with this observation, while being similar in sequence to the contryphan-Vc1 toxin and the other two doppelganger toxins identified here, the matching signaling DREP family shares very little similarity with actual contryphan toxins. The unusual evolution of this doppelganger toxin family will be addressed in more detail below.

We note that homology searching identified unknown signaling peptides in both mollusks and annelids but did not detect any novel signaling peptides in zebrafish. Possible explanations for this are provided in the discussion.

In addition to the five new families, we were able to identify eight previously described doppelganger conotoxin/signaling peptide pairs: conopressins (Cruz et al. 1987), conoRFamide (Maillo et al. 2002), conoCAP (Möller et al. 2010), cono-neuropeptide F/Y (Wu et al. 2010), con-insulins (Safavi-Hemami et al. 2015), elevenin (Robinson et al. 2017a), prohormone 4 (Robinson et al. 2017a), and consomatin (Ramiro et al. 2022) but did not detect conoMAP (Dutertre et al. 2006) or contulakin-G (Craig et al. 1999). ConoMAP shares sequence similarity with signaling peptides of the myoactive tetradecapeptide family (MATPs). ConoMAP was only described in a single species of worm hunter, *Conus vitulinus*, whose transcriptome has not been made available yet and was therefore not included. The other 43 species analyzed here do not appear to express toxins with significant sequence similarity to MATPs. Contulakin-G was identified from a single species of fish hunter, *Conus geographus*. Although the transcriptome of *C. geographus* containing the contulakin-G sequence was included here, the toxin only shares limited sequence similarity with the fish neurotensin precursor (*i*.*e*., 4 identical amino acids) escaping the homology search applied.

### Doppelganger toxins and their DREPs share a high degree of similarity in the peptide region

While doppelganger toxins evolved to mimic the signaling peptides of their target organism, the non-toxin-encoding regions of the precursors are presumably under little if any evolutionary pressure to mimic the signal sequence or the pro-region(s) of the prey precursor. Thus, we hypothesize that the precursors of doppelganger toxins and their DREPs may only show significant similarity in the region that encodes the mature peptide. To investigate this, we aligned each of the toxins to their respective DREPs and quantified the number of identical amino acids in the signal sequence, the peptide region, and the spacer region(s). Indeed, we found that the toxins are significantly more similar to the prey protein in the peptide region compared to the signal sequence and the spacer regions (where applicable) (**Supplementary Figure 2)**. Due to the low number of toxins for Hairpin DREPs (n=3), we were not able to statistically quantify the amino acid percentage identity in the different regions. However, in the other four cases, there is a clear trend toward higher similarity in the peptide region. Overall, we find that this region displays between 35 - 55 % identity compared to only 12 - 28 % for the signal sequences and spacer regions.

### Structural predictions further suggest that doppelganger toxins target prey signaling peptide systems

Signaling peptide action is mediated via binding to membrane proteins, mostly belonging to the GPCR family. Such binding is contingent on the complementarity of the receptor ligand-binding site and the tertiary structure of the peptide ligand. Whereas many signaling peptides are disordered in solution, several have a well-defined structure; in particular, those that contain disulfide bonds. It can be hypothesized that most doppelganger toxins conserve their overall three-dimensional structure through evolution to preserve the ability to stimulate the prey target receptor although some structural features may adapt for improved function (e.g., higher stability, better selectivity). To investigate this, we obtained structural predictions of doppelganger toxins and their prey DREPs using AlphaFold2 (Jumper et al. 2021).

For the Triangle DREP and toxin pair, we performed a structural prediction of the *Conus rattus* doppelganger toxin and *P. dumerilii* DREP peptide, representing the most highly expressed toxin and the founding member of the Triangle DREP family, respectively. Both predicted structures (average pLDDT 72.4 and 67.6, respectively) form three alpha-helices in a triangular loop linked by a disulfide bond with disordered N- and C-termini (**Figure 3A**). Even though the sequences show less than 50 % sequence identity, the predicted structures are almost identical (rmsd 0.71). We note that the amino acid residues of the toxin that show the highest degree of evolutionary divergence (as measured by rate4site score) are primarily located on the exterior of the structure. These sites are likely interacting with the binding sites on the target receptor and have specifically been modified to provide novel functions or selectivity profiles of the toxins compared to the endogenous ligands.

**Figure 3:**
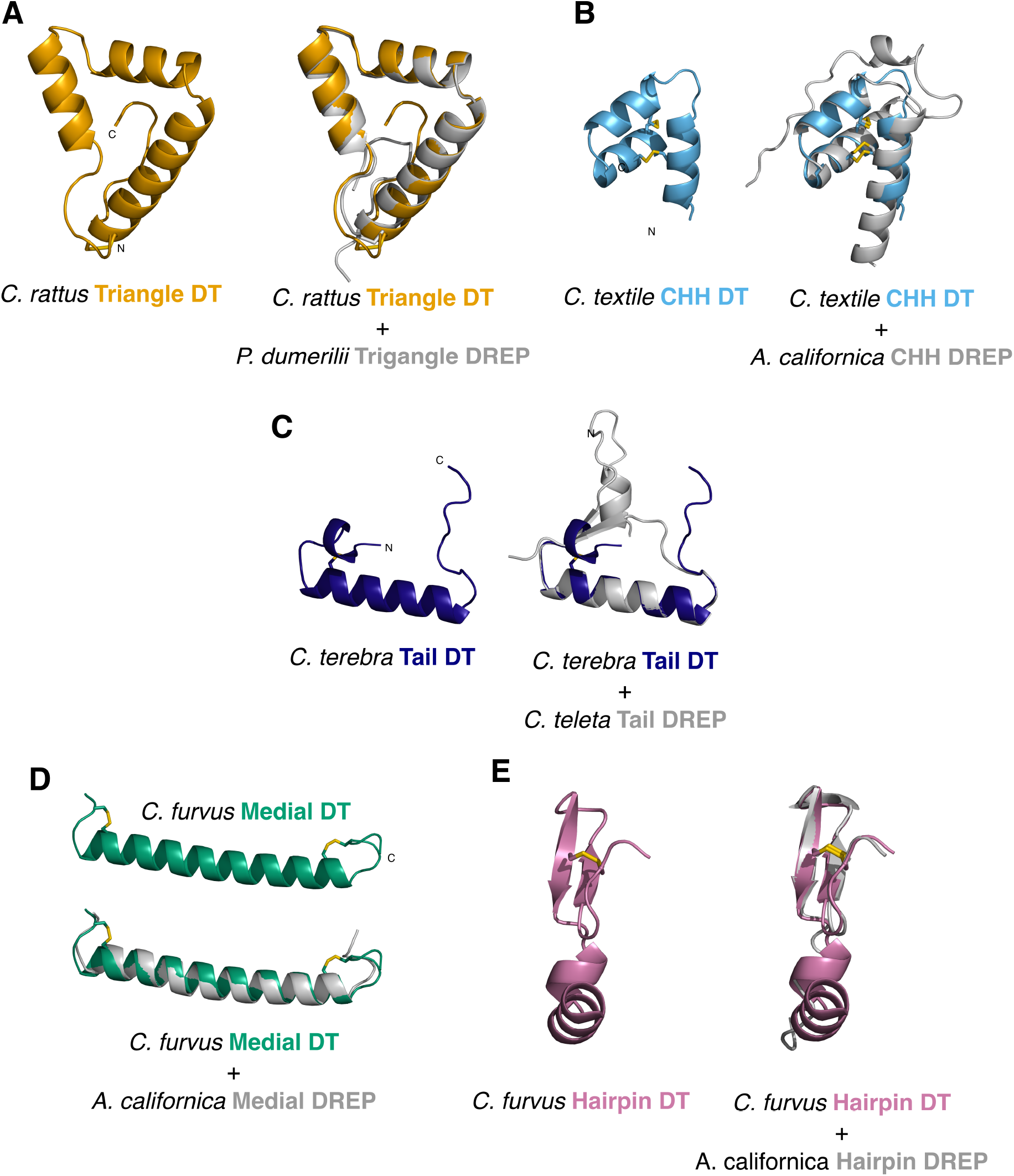
Structural predictions (Alphafold2) and alignments of doppelganger toxins and DREPs highlight their structural similarity despite limited sequence identity. (**A**) *Conus rattus* Triangle doppelganger toxin 1 (left) and alignment with *Platynereis dumerilii* Triangle DREP (right). (**B**) *Conus textile* CHH doppelganger toxin (left) and alignment with *Aplysia californica* CHH DREP (right). (**C**) *Conus terebra* Tail doppelganger toxin (left) and alignment with *Capitella teleta* Tail DREP (right). (**D**) *Conus furvus* Medial doppelganger toxin (top) and alignment with *A. californica* Medial DREP (bottom). (**E**) *C. furvus* Hairpin doppelganger toxin (left) and alignment with *A. californica* Hairpin DREP (right).

Structural predictions of the CHH DREP and toxin pair suggest that both the highly expressed *C. textile* toxin and the Aplysia DREP peptide fold to form disulfide bonds between cysteines I-III and II-IV (**Figure 3B**). Both predicted structures have three alpha-helices, and the *A. californica* peptide has an additional short helix in the flexible N-terminus (average pLDDT 91.7 and 67.0). Structural alignment results in a good overlay of the orderly-folded regions (1.1 rmsd). The evolutionarily diverging residues of the toxins are mostly facing outward and are in helix 2 and around the interface region between helix 1 and 3, suggesting that receptor interactions are likely to take place in these regions of the peptide.

The predicted structure of the Tail doppelganger toxin from *Conus terebra* (average pLDDT = 69.1) has a single alpha-helical region located C-terminally of the single cysteine loop (**Figure 3C**). The matching Tail DREP structure from the annelid *C. teleta* (average pLDDT = 67.14) also shows a single alpha-helical region following the cysteine loop and contains a short segment of parallel beta-sheets toward the N- and C-termini. An alignment of the two structures gives an rmsd of 0.7.

Structural predictions of the Medial DREP peptide and toxin pair suggest a long alpha-helical fold (Aplysia DREP: average pLDDT = 61.82, *C. furvus* toxins: average pLDDT= 61.76). The confidence of these structures is low, and it is likely that these toxins and signaling peptides exist in a disordered state when unbound (low pLDDT-values have been linked to protein disorder (Tunyasuvunakool et al. 2021)) or that these sequences are not feasible for Alphafold-based structural predictions (**Figure 3D**).

Predictions of the structures of the Hairpin doppelganger toxin from *C. furvus* and Hairpin DREP from Aplysia yielded average pLDDT of 70.86 and 55.1, respectively (using the first peptide copy of the Aplysia precursor). While the confidence for the Aplysia structure is low, the two peptides align well with 1.01 rmsd (**Figure 3E**). The structure of contryphan-Vc1 from the Hairpin doppelganger family has been experimentally determined (PDB:2N24) (Robinson et al. 2016) and aligns well with the predicted structure of the *C. furvus* toxin (1.83 rmsd). These structures are all characterized by a single disulfide bridge-directed beta-hairpin that connects the N-terminal region of the peptides with the second of two anti-parallel beta-strands. As demonstrated for contryphan-Vc1, this structure provides a substantial thermal stability (Robinson et al. 2016). Our findings on the predicted structure of the Aplysia Hairpin DREP peptide suggest that this stable fold may have already existed in the signaling peptide family that gave rise to the toxins.

Collectively, the high similarity between the predicted structures of DREPs and their corresponding doppelganger toxins further suggests that the toxins specifically mimic the herein identified signaling peptides thereby likely disrupting important endogenous signaling events in prey.

### Structural predictions of CHH DREP identify the first spiralian member of the CHH hormone family

Protein three-dimensional structures are believed to be more conserved than the corresponding amino acid sequences (Illergård et al. 2009). We, therefore, tested if any of the predicted structures of the doppelganger toxins or their DREPs showed resemblances to known peptides with experimentally verified structures by using the structural similarity search method DALI (Holm 2022). This could provide further insight into the function and evolutionary origin of the identified peptides.

Whereas most of the searches only resulted in low similarity hits, a search for structural homologs of the *Aplysia* CHH DREP peptide yielded several close matches. The top hits were structures of k-Ssm1a (PDB: 2M35) and Ssd609 (PDB: 2MVT) toxins from the centipede *Scolopendra subspinipides*, the insecticidal toxin Ta1a from the funnel spider *Eratigena agrestic* (PDB: 2KSL), and notably the structures of crustacean hyperglycemic hormone (CHH) (PDB: 5B5I) of the kuruma prawn, *Panaeus japonicus*. We observe a strong structural resemblance between the predicted structure of the Aplysia peptide to both the k-Ssm1a and Ssd609 toxins (2.1 and 1.9 rmsd) and the kuruma prawn CHH (2.7 rmsd), even though the sequences only share 26 %, 19 %, and 14 % sequence identity, respectively (**Figure 4**). The structural similarity strongly suggests that the herein-identified Hairpin DREP sequences encode signaling hormones belonging to the CHH superfamily. This would be the first example of a signaling peptide belonging to the CHH family found outside of Ecdysozoa, a heterologous group of protostomes that includes nematodes and arthropods. Another interesting feature is that the molluscan DREP peptides and their doppelganger toxin pairs only contain two disulfide loops, whereas both the arthropod toxins and arthropod hormones contain three.

**Figure 4:**
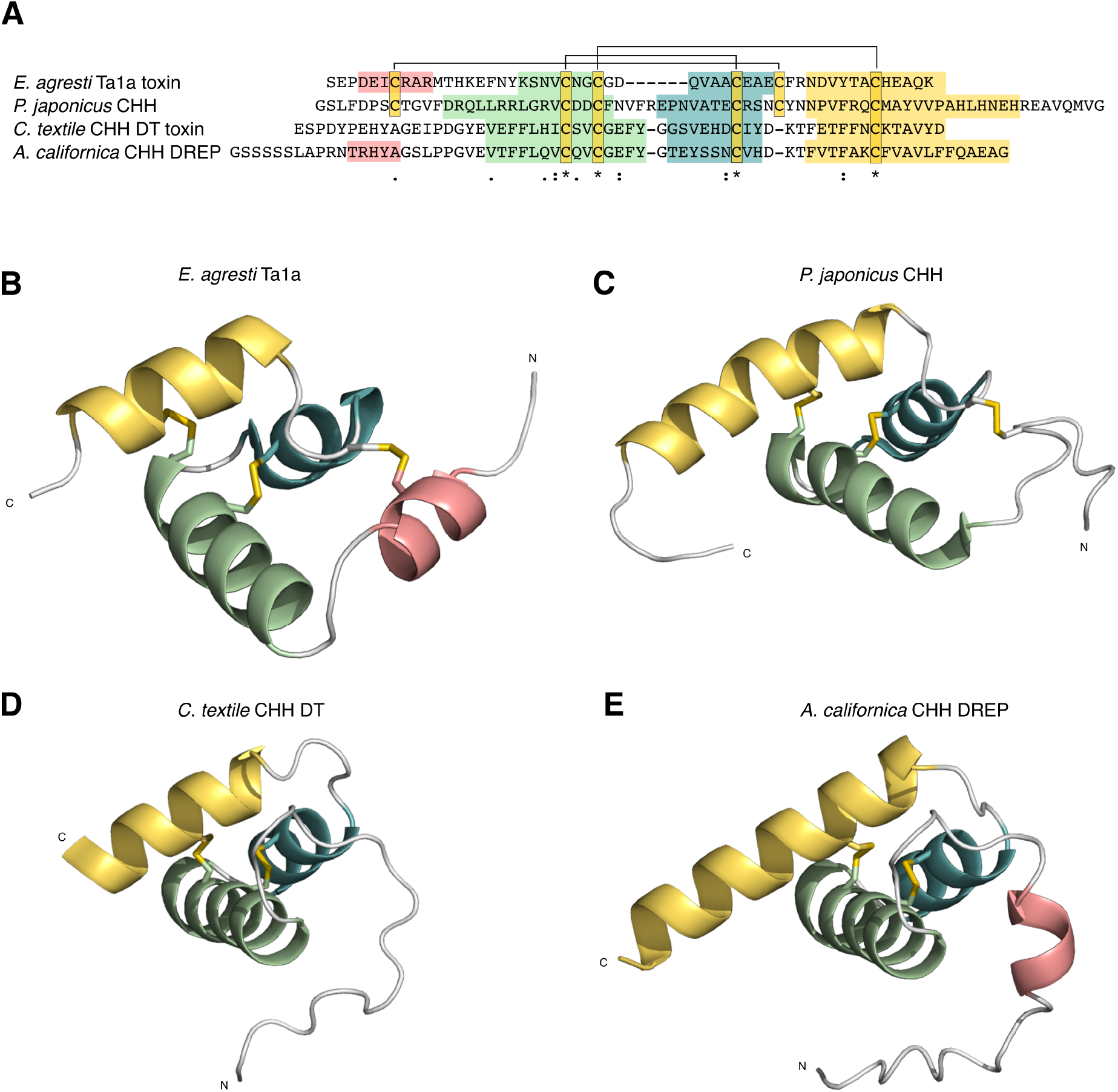
Structural similarity suggests that the CHH DREP family is related to the crustacean hyperglycemic hormone (CHH)-superfamily of signaling peptides. (**A**) Multiple sequence alignment of funnel spider (*Eratigena agrestic*) Ta1a CHH-toxin, *Panaeus japonicus* CHH, *Conus textile* CHH doppelganger toxin, and *Aplysia californica* CHH DREP show limited sequence similarity and share only two out of three disulfide loops. Coloration corresponds to alpha helices shown in B-E. (**B**) Ta1a toxin from funnel spider *E. agresti* (PDB: 2KSL), (**C**) *P. japonicus* CHH (PDB: 5B5I), (**D**) Alphafold2 structural prediction of *C. textile* CHH doppelganger toxin, (**E**) Alphafold2 structural prediction of *A. californica* CHH DREP. (B-D) have similar tertiary structures.

### Doppelganger toxins are highly expressed in the venom gland

We next investigated the expression levels of the identified doppelganger toxins in the respective venom gland datasets. Conotoxin expression is highly variable with expression levels generally ranging between 10 - 100,000 transcripts per million (TPM) (Phuong et al. 2016; Robinson et al. 2017b). While a very low level of expression can be indicative of contamination by surrounding tissues or cross-contamination from multiplexing, highly expressed transcripts almost certainly encode toxins. For all five doppelganger families, we found at least one highly expressed transcript (>1,000 TPM), supporting that these families are secreted toxins and are functionally important in at least some cone snail species (**Supplementary Figure 3)**. Additionally, none of these doppelganger toxins were expressed in cone snail neuroendocrine tissue, salivary glands, or the foot, further suggesting that these peptides are specific components of the venom.

### Tissue-specific transcriptomes confirm the expression of DREPs in neuroendocrine and other secretory tissues of mollusks and annelids

If the identified DREPs encode signaling peptides, we would expect that at least some of these transcripts are primarily expressed in neuroendocrine and/or other secretory tissues. To test this, we quantified the expression of DREPs in tissue-specific transcriptomes of *A. californica* (generated here) and publicly available datasets of the mollusk *Doryteuthis pealeii* (longfin inshore squid) and annelid *Lumbricus rubellus* (red earthworm). Tissue-specific datasets were not available for the two model annelids, *C. teleta* and *P. dumerilli*.

Using the *Aplysia* datasets, we did not identify orthologs of Triangle DREP or Tail DREP. *Aplysia* CHH DREP is, on the other hand, expressed in all eight sequenced ganglia (especially in the abdominal ganglion) and nerves but missing in non-neuronal transcriptomes (**Supplementary Figure 4A**). We found that *Aplysia* Medial DREP is expressed at relatively low levels in the pleural ganglion (9.23 TPM) but not in non-neuronal tissues. Lastly, *Aplysia* Hairpin DREP is expressed in all the neuronal transcriptomes, particularly in the pedal ganglion, and also highly expressed in the salivary gland. We also observe low expression in the foot. The salivary glands are secretory organs that are known to express some signaling peptides (Dawidson et al. 1997). However, it is also possible that Hairpin DREP serves a non-signaling-related function in the saliva gland and potentially the foot of *A. californica*.

In the squid transcriptomes, we found orthologs of Triangle, CHH, and Hairpin DREPs in various tissues. Triangle DREP is highly expressed in the brain, nerves, brachial, vertical, and optical lobes, and to a lower degree in the testes and buccal mass. CHH DREP is expressed in the brain and brachial lobe, and Hairpin DREP expression was detected in several neuronal tissues (brain, brachial lobe, the giant fiber, vertical lobe, nerves, optical lobe, retina, and stellate ganglion), but also in other, non-neuronal but secretory tissues (buccal mass, hepatopancreas, kidney, and skin) (**Supplementary Figure 4B, supplementary File 6)**.

Using the earthworm transcriptomes, we found evidence of expression of Triangle and Hairpin DREPs in the nerve cord and neural ganglion. CHH DREP is expressed in transcriptomes from the body wall and the clitellum (the non-segmented, band-like region of earthworms with glandular cells), and Tail DREP in the calciferous (calcium secreting gland), chlorogog, crop, and gizzard (digestive organs) (**Supplementary Figure 4C, Supplementary File 6)**. Medial DREP could be retrieved from other annelid datasets (shown below) but was not expressed in any of the queried tissue-specific transcriptomes from *L. rubellus*. The expression of Medial DREP may be restricted to specific cell populations or developmental stages that were not captured in the available datasets. Nonetheless, our combined findings from *A. californica, D. pealeii*, and *L. rubellus* show that the identified DREPs are functionally expressed and predominantly associated with neuroendocrine and secretory tissues.

### Doppelganger-related peptides are widely present in mollusks and annelids

Having established the expression of DREPs in several transcriptomes, we searched for additional DREP orthologs in mollusks, annelids, and other metazoans. We found that all five families are present in other mollusks and annelids (**Supporting files 6, Figure 5**). These precursors are all secreted, have similar precursor architecture (*e*.*g*., location of the predicted peptide), have an identical number of cysteines, and similar processing sites as the toxins and the initial prey sequence. Using a more sensitive psi-BLAST strategy, we furthermore identified multiple genes encoding Triangle DREP-like peptides in other protostome phyla, including Rotifera, Arthropoda, Platyhelminthes, and Tardigrada (**Figure 5, Supplementary File 7)**. These precursors all show the same architecture as the other Triangle DREP peptides from mollusks and annelids with peptides located immediately after the signal sequence and terminates 4 - 10 amino acids downstream of the long disulfide loop in a classical processing site. In CLANS clustering analyses the precursors from rotifers, arthropods, platyhelminths, and tardigrades cluster with the molluscan and annelid sequences (**Supplementary Figure 5)**. In addition to the significant sequence similarity, we also found that each of the five DREP families shares common features of intron position and phase across phyla (**Supplementary File 8**), corroborating that the identified orthologs indeed belong to the same family as the initial prey sequence.

**Figure 5:**
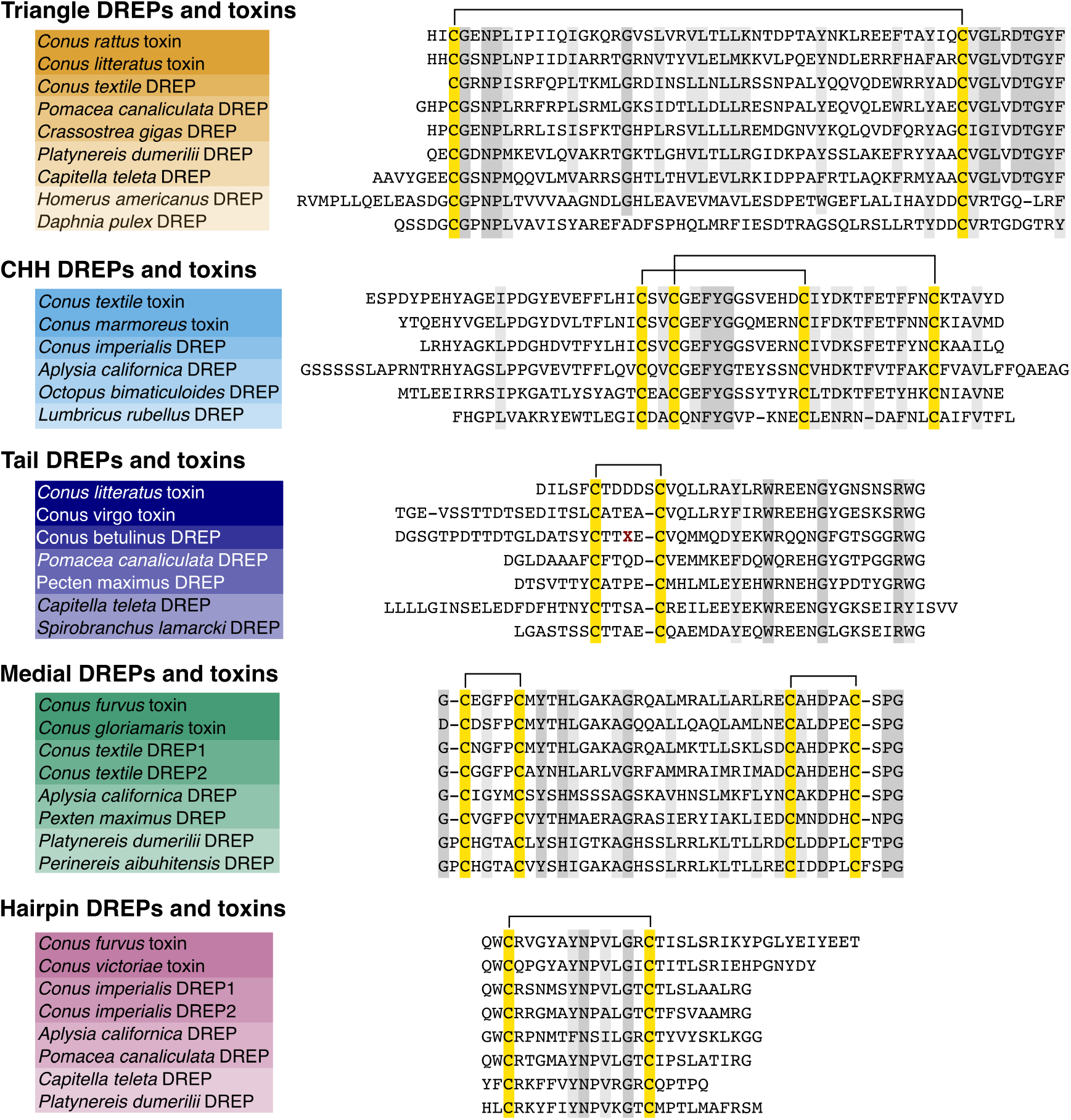
Multiple sequence alignment of representative mature toxins and signaling peptides of the five doppelganger toxin and DREP families. Alignments highlight high sequence similarity of the toxins and DREPs, including conserved cysteine scaffolds. Two endogenous Medial and Hairpin DREPs are found in cone snails (DREP1 and 2). The *Conus betulinus* Tail DREP is only a partial sequence with a sequencing error (dark red X).

Furthermore, with two exons separated by a phase 2 intron, the CHH DREP identified here mirrors the proposed gene structure of the ancestral ecdysozoan CHH gene (Montagné et al. 2010), supporting that the CHH DREPs belong to the CHH superfamily.

### Doppelganger toxins likely evolved from orthologs of the signaling genes they mimic

To investigate if the doppelganger toxins identified here evolved through the recruitment of endogenous signaling genes into the venom, we queried the transcriptomes of the circumoesophageal nerve ring from several cone snail species for sequences that could have given rise to the doppelganger toxins. The circumoesophageal nerve ring is a structure of cone snail nerve ganglia (the cerebral, pedal, and pleural ganglia) known to express signaling peptide genes, some of which are known to have been recruited into the venom gland (Koch et al. 2022; Safavi-Hemami et al. 2016).

In all cases we were able to recover homologous transcripts from the nerve rings of *Conus textile, Conus rolani*, and/or *Conus imperialis*, or from the genomes of *Conus ventricosus* or *Conus betulinus* (**Figure 5, Supplementary File 9)**. However, we were only able to identify a partial sequence of the likely endogenous cone snail Tail DREPs. In two cases (Medial DREPs and Hairpin DREPs) we identified two paralogous genes in the nerve rings (ancestral gastropods are believed to have undergone a whole-genome duplication, and this event has led to many pairs of paralogs in present-day cone snails (Pardos-Blas et al. 2021)).

The endogenous genes retrieved here show common characteristics of signaling peptide precursors such as the presence of an N-terminal signal sequence and proteolytic cleavage sites flanking the signaling peptide-encoding region. They also show the same features of intron positions and phases identified in other molluscan and annelid DREPs identified above (**Supplementary file 8**). Multiple sequence alignment of the nerve ring precursors with the corresponding toxins clearly shows that these proteins are related (**Figure 5**), and the observed sequence similarity even extends into the 3’ and 5’ untranslated regions (**Supplementary Figure 6)**.

**Figure 6:**
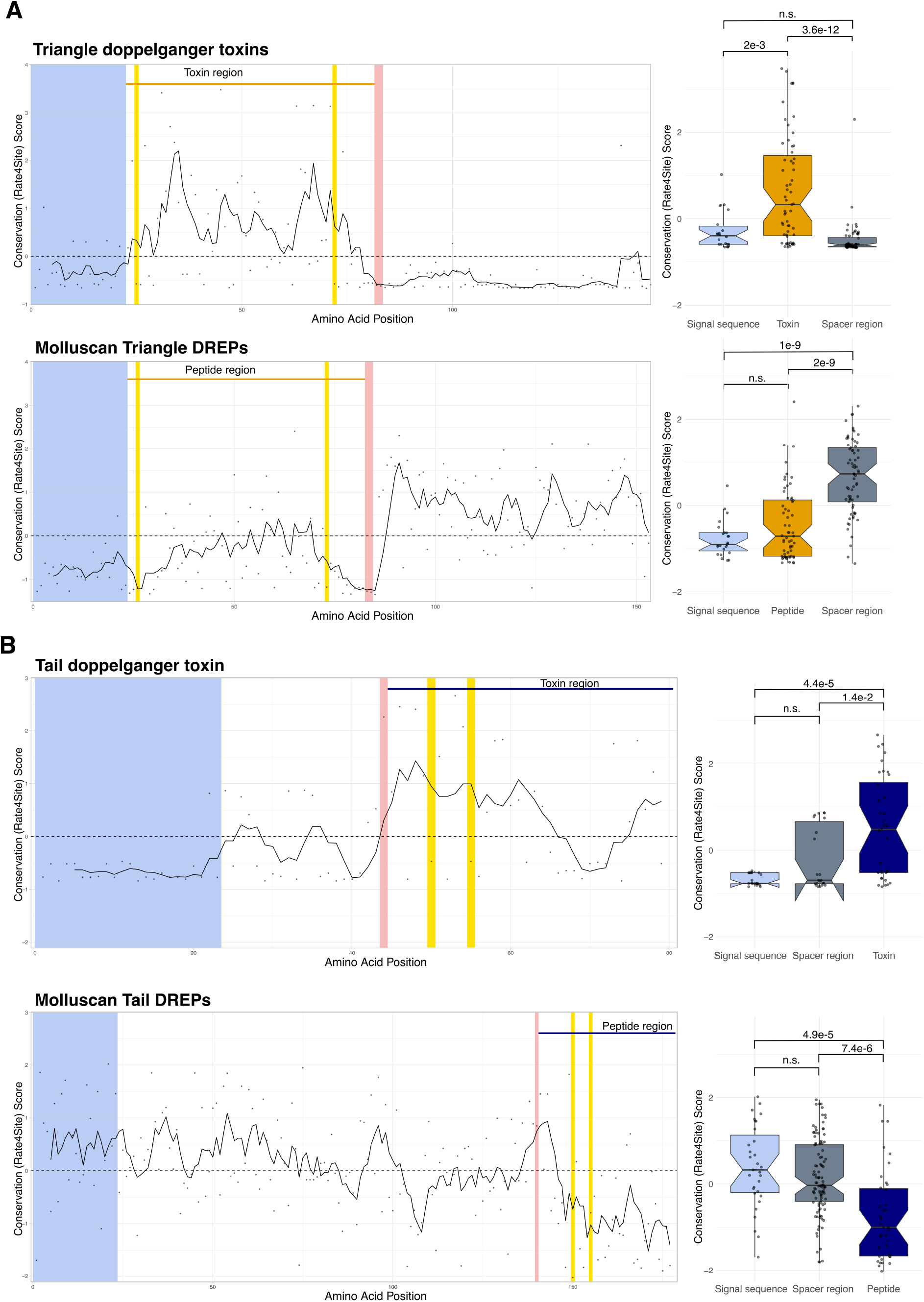
Evolutionary trace analyses show different conservation (rate4site) scores in the toxin/peptide regions compared to the signal sequence and spacer region(s). (**A**) Position-specific rate4site scores for Triangle doppelganger toxin represented by *Conus litteratus* toxin and molluscan Triangle DREPs represented by the endogenous *Conus textile* precursor. Wilcoxon rank-sum test shows significant differences between the toxin region compared to the signal sequence and spacer region. (**B**) Position-specific rate4site scores for Tail doppelganger toxin represented by *Conus terebra* toxin and molluscan Triangle DREP represented by the endogenous *Pomacea canaliculata* precursor. Wilcoxon rank-sum test shows significant differences between the toxin region compared to the signal sequence and spacer region (p values shown). The signal peptide is depicted in light blue, processing sites are in red, and cysteines in yellow. The peptide and toxin regions are shown above the graphs. Spacer regions are defined as the non-signal sequence/peptide/processing site regions.

To gather more evidence for the evolutionary relatedness of doppelganger toxins and their DREPs we mined the genomes of the two worm-hunting species *C. ventricosus* and *C. betulinus* for doppelganger toxin-encoding sequences. As of writing, the genomes of snail-hunters have not been sequenced or made available yet. Using the worm-hunter genomes, we were able to retrieve sequences of the two gene families that were identified in the venom glands of worm-hunting species, Triangle and Tail doppelganger toxins, but not those that were exclusively expressed in snail hunters (*i*.*e*., CHH, Medial, and Hairpin doppelgangers). We found that the gene structure of Triangle doppelganger toxins mirrors that of the endogenous signaling peptide, both of which have a single phase 1 intron located within the disulfide loop (**Supplementary File 8**).

Tail doppelganger toxins are located on a single exon. Unfortunately, this gene structure could not be compared to that of endogenous Tail DREP, as we could only recover a partial sequence of the endogenous cone snail DREP gene.

While we could not perform additional gene structure analyses for the remaining doppelganger families, the sequence similarities observed at the transcript level strongly suggest that the identified doppelganger toxins evolved from the highly conserved nerve ring proteins identified here. This further supports the hypothesis that, on multiple occasions, endogenous signaling peptide-encoding genes were recruited into the venom gland where they diversified and neofunctionalized to specifically target the related peptide signaling system of prey.

### Evolutionary trace analyses show contrasting patterns of conservation in doppelganger toxins and DREPs

Using a single doppelganger toxin/DREP pair we previously showed that signaling peptides and peptide toxins experience contrasting selection pressures that reflect their divergent biological functions (Koch et al. 2022). Having identified five new families of doppelganger toxins and their DREPs allowed us to comprehensively investigate differential rates of evolution between the toxins and their signaling peptide pairs using evolutionary trace analyses. Such analyses identify the relative level of conservation for each site in multiple sequence alignments of related sequences. Whereas signaling peptide precursors show a higher level of conservation in the mature peptide region compared to the signal and spacer regions, toxin precursors show an elevated level of amino acid diversity in the toxin region in conjunction with a few highly conserved residues (Fry et al. 2009; Woodward et al. 1990). Here, we performed evolutionary trace analysis to investigate if the patterns of evolution of the identified doppelganger toxins and their DREPs matched those seen for other toxin and signaling peptide precursors, respectively.

Evolutionary trace analysis of Triangle doppelganger toxins confirmed the presence of mostly divergent sites within the toxin regions in conjunction with a few conserved residues including the cysteine residues (**Figure 6A**). The C-terminal spacer peptide is, on the other hand, highly conserved. The corrected Wilcoxon rank-sum test (commonly referred to as the non-parametric version of the t-test, not assuming a normal distribution) also shows significantly different rate4site scores in the peptide, signal sequence, and spacer region (**Figure 6A**). In contrast, we find that the peptide region of the molluscan and annelid Triangle DREP signaling peptides shows an elevated level of conservation compared to the spacer region, suggesting that these precursors indeed encode a conserved signaling peptide (**Figure 6A, Supplementary Figure 7**). It has been suggested that the distinct rates of evolution in the toxin, spacer, and signal sequence of conotoxins are due to these regions being located on individual exons. When we compared the evolutionary trace of the two Triangle doppelganger exons, we did observe a significant difference between the first and second exons (p = 4.1e-4). However, this difference was not as pronounced as when comparing the functional regions (i.e. the signal sequence, toxin region, and spacer region) (**Supplementary Figure 8**).

**Figure 7:**
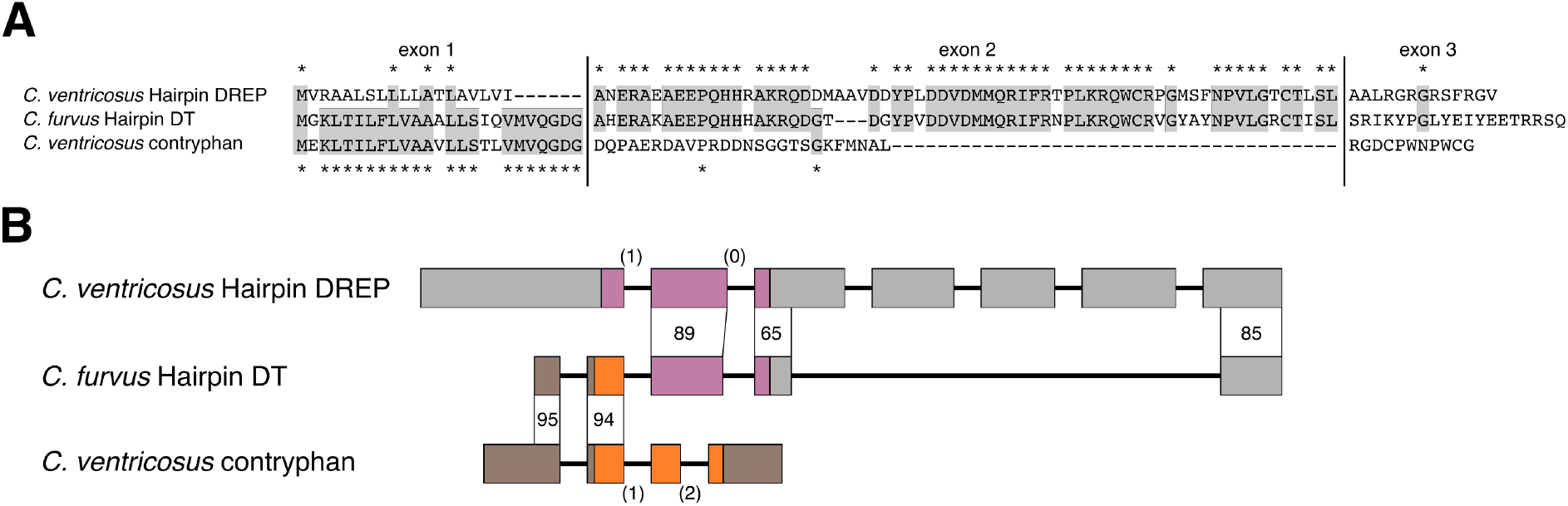
Hairpin doppelganger toxins evolved by exon shuffling of the cone snail endogenous Hairpin DREP and contryphans. (A) Multiple sequence alignment of the amino acid sequences shows high similarity of Hairpin doppelganger toxin to contryphan/O2 toxin in the signal sequence region located on the first coding exon, and a high similarity to cone snail endogenous Hairpin DREP in the second exon encoding the mature toxin. (B) Genes of *Conus ventricosus* Hairpin DREP and contryphan are consistent with an origin of Hairpin doppelganger toxin by exon shuffling. The *C. ventricosus* Hairpin DREP gene consists of 7 exons (wide boxes) with the open reading frame located on exons 1-3 (purple; UTR: gray) separated by a phase 1 and phase 0 intron (shown in parantheses). The *C. ventricosus* contryphan gene consists of 4 exons with the open reading frame located on exons 2-4 (orange) separated by a phase 1 and phase 2 intron. The *C. furvus* doppelganger toxin gene shares high identity with the 3’ UTR region of contryphan exon 1 and exon 2 (95 and 94 %), and high identity with Hairpin DREP exon 2, 5’ end of exon 3, and 5’ end of exon 7 (89, 65, 85 %).

Consistent with findings for Triangle doppelganger toxins, Tail doppelgangers show a significantly more diverging mature toxin region compared to the spacer region (**Figure 6B**). On the other hand, the molluscan and annelid Tail DREP precursors, show the opposite pattern with more conserved peptide regions when compared to the spacer region (**Figure 6B, Supplementary figure 7**).

Comparison of the evolutionary conservation between the mature toxin/peptide region and the spacer region of the CHH doppelganger/DREP families was, unfortunately, not possible, as the mature peptides span the entire precursor following the signal sequence with an apparent lack of a spacer region. However, while we could not analyze any spacer regions, when we analyzed the precursors lacking the N-terminal signal sequence, we do observe higher sequence variability of the toxins compared to the signaling peptides (**Supplementary Figure 7**).

When we examined the evolutionary trace of the Medial doppelganger toxins, we observed a wide distribution of evolutionary conservation within the predicted toxin region: Some amino acid residues are highly conserved, such as the cysteine residues, whereas others, particularly in the central region, are variable. The amino acids of the spacer regions are, on average, less conserved than those of the mature toxin region. Correspondingly, there is no significant difference between the overall rate4site score of the peptide and spacer regions. The molluscan Medial DREP precursors show the expected evolutionary trace of neuroendocrine signaling peptides, where the peptide region is more conserved than both the N- and C-terminal spacer regions (**Supplementary Figure 7**).

Unfortunately, we only identified three Hairpin doppelganger toxins making evolutionary trace analysis less informative for this family. With such a low number of examples, it is harder to estimate residue conservation and the observed conservation might be more reflective of phylogenetic sampling rather than true conservation. Nonetheless, we did observe a higher number of diverging amino acids in the predicted toxin region compared to the spacer region (**Supplementary Figure 7)**. Having a larger number of the endogenous molluscan and annelid Hairpin DREP counterparts, we do find elevated conservation of the mature peptide region of these peptides (p = 5.5e-5 and p = 2e-8 for mollusks and annelids, respectively, **Supplementary Figure 7**).

Despite some differences in the evolutionary trace analyses for the five doppelganger toxins and DREP families, we generally observe contrasting patterns of evolution between the toxin and signaling peptide sequences. The mature toxin regions are, on average, more divergent and have a higher deviation than the surrounding spacer regions. In contrast, the signaling precursors are generally conserved in the peptide-encoding region. This is consistent with diversification and neofunctionalization of toxin genes following recruitment from a conserved endogenous signaling gene into the venom gland.

### Hairpin toxins evolved through exon shuffling

The underlying molecular mechanism that results in conotoxin diversity is not fully understood. While differential rates of evolution in the distinct functional units of conotoxin precursors play an important role, other mechanisms have also been proposed (Pi et al. 2006). We noticed that the signal sequence of Hairpin doppelganger toxins belongs to the Contryphan/O2 toxin superfamily, but the remaining regions are very distinct from other sequences belonging to the contryphan/O2 toxin superfamily. Furthermore, the Hairpin doppelganger toxins share high sequence similarity with the endogenous cone snail Hairpin DREP family, but only in the region containing the spacer regions and mature peptides (**Figure 7A**). Based on these observations, we hypothesized that Hairpin doppelganger toxins evolved by exon shuffling to create a Contryphan/O2-Hairpin DREP chimera. Exon shuffling has been observed in other venomous animals (Wang et al. 2016).

To investigate this hypothesis, we annotated the endogenous Hairpin DREP and Contryphan/O2 toxin genes in the genome of *C. ventricosus* based on transcripts retrieved from *C. rolani* Hairpin DREP and the *C. boavistensis* Contryphan toxin (**Supplementary File 10)**. We found that the Hairpin DREP gene consists of 7 exons with the Hairpin DREP precursor located on exons 1 - 3 that are separated by a phase 1 and a phase 0 intron (**Figure 7B**). The *C. ventricosus* contryphan gene consists of 4 exons with the venom precursor located on exons 2 - 4 that are separated by a phase 1 and a phase 2 intron (**Figure 7B)**. When we aligned the Hairpin doppelganger toxin gene from *C. furvus* to the *C. ventricosus* contryphan gene, we found that the 5’ UTR and the region encoding the signal sequence aligns with 95 % identity, but that the remaining 3’ end of the *C. furvus* gene and 3’ UTR only aligns with 28 % identity (**Supplementary File 11**). Conversely, when the *C. furvus* Hairpin toxin gene is aligned to the *C. ventricosus* Hairpin DREP gene, we only observe 18 % identity in the 5’ UTR and region encoding the signal sequence, whereas the remaining 3’ end of the gene and 3’ UTR aligns locally with 65 – 89 % identity (**Supplementary File 11**). The most parsimonious explanation is that the Hairpin doppelganger toxins evolved by shuffling of exons 1 and 2 of a contryphans gene with exons 2, 3, and 7 of the endogenous Hairpin DREP gene; a fusion made possible by the two phase-1 introns in both the Hairpin DREP and contryphan genes. Because the contryphan/O2 superfamily is found throughout Conus (Grant et al. 2004; Jimenéz et al. 1996; Massilia et al. 2001), these toxins are most likely evolutionarily older than Hairpin doppelganger toxins, which seemingly are confined to snail hunters. This leads us to believe that the Hairpin doppelganger toxins adopted the contryphan/O2 signal sequence rather than the other way round.

## Discussion

Signaling peptides are essential to animal biology, but *de novo* discovery is often difficult. In this study, we used a method centered around doppelganger toxins from venomous marine cone snails to discover putative signaling peptides of the prey organisms. Using this approach, we identified five undescribed molluscan and annelid protein families that we propose encode novel signaling peptides and their related doppelganger toxins.

Several lines of evidence suggest that the doppelganger toxins we discovered in cone snail venom gland transcriptomes are in fact real toxins that define new toxin gene superfamilies. First, all the sequences contain N-terminal signal sequences that target the peptides to the secretory pathways and are used to classify toxins into distinct superfamilies. Second, at least one member of each superfamily is highly expressed in the venom gland and not in other tissues. Third, we find the toxin sequences display conserved signal and spacer regions combined with hypervariable toxin regions, a well-described feature of cone snail toxins. Forth, we recovered the conserved nerve ring genes that gave rise to the toxins. Finally, we demonstrate the presence of an emerging characteristic pattern of contrasting evolutionary conservation between doppelganger toxins and the DREPs they originated from (**Figure 5)**. Collectively, these findings leave little doubt that the herein identified doppelganger toxins are *de facto* conotoxins. Future studies using tandem mass spectrometric (MS/MS) sequencing will be needed to confirm the presence of the translated peptides in venom. These studies will also be important for determining the presence of post-translational modifications (commonly found in conotoxins) and alternative proteolytic processing sites (Buczek et al. 2005).

Similarly, based on multiple lines of evidence, we propose that the identified DREP families encode previously unknown signaling peptides. First, all members of these families contain an N-terminal signal sequence, showing that they encode secreted proteins. Second, we identified enzymatic processing sites characteristic of signaling peptides (basic and dibasic amino acids). Third, evolutionary trace analyses show a pattern of conservation characteristic of signaling peptides (but contrasting to toxins). Forth, all five families are found throughout different classes of Mollusca and Annelida, and in one case we also identified orthologs in several other protostome phyla. Fifth, we observed expression of these genes in cone snail nerve ring tissue and, in most cases, also in neuroendocrine and/or secretory tissues of the mollusks *A. californica* and *D. pealeii*, and the annelid *L. rubellus*. Jointly, these findings strongly support the hypothesis that these families encode new bilaterian signaling peptides.

As for the doppelganger toxins, future studies using MS/MS sequencing will be needed to confirm the presence of the translated peptides, including their potential modifications. Our findings on the tissue-specific expression of these peptides could inform on the organism and tissue to be used as a starting point for MS/MS-based detection.

We have gathered strong evidence that the new prey protein families encode neuroendocrine signaling peptides of biological importance. However, in the absence of functional data, it cannot be ruled out that the peptides discovered here have alternative functions. Several venomous animals express protease inhibitors, which are typically 50-60 amino acids in length and processed from a secreted precursor (Mourão and Schwartz 2013). Most peptides identified here are shorter, and it is thus unlikely, albeit not impossible, that these are protease inhibitors. Alternatively, the sequences identified here could encode other enzyme inhibitors (e.g., kinase inhibitors), proteins with alternative functions (e.g., pore-forming or antimicrobial function), or proteins of yet unknown function. Establishing that these peptides are indeed signaling peptides will require future functional studies, ideally combined with the identification of their molecular targets. However, if, as we propose, these sequences encode signaling peptides, these peptides and the systems they regulate are likely of functional importance in prey. The evolutionary cost of producing toxins is high, and toxins that target systems of little importance should, in principle, be selected against. We consequently suspect that the signaling peptides identified here regulate critical functions in mollusks, annelids, and other organisms and will likely be of importance to future neurobiological research.

Despite the lack of functional information, the new doppelganger/DREP pairs have already revealed several new insights into peptide evolution and putative function.

For example, using structural homology searches we observed that one of the new DREP peptide families showed significant similarity to CHH peptides found in arthropods and nematodes – peptides that have been firmly established as signaling peptides (Chen et al. 2020). CHH was originally defined by its hyperglycemic activity (Abramowitz et al. 1944). However, it has become clear that CHH and its related peptides have a wide range of physiological functions in metabolism, water and ion balance, development, immune regulation, molting, and ovarian maturation (Chen et al. 2020). When we compared the gene structures (i.e., number, phases, and positions of introns) of the CHH DREP genes with those encoding arthropod CHH, we found identical patterns serving as evidence for the common ancestry of these signaling peptides. This finding on the existence of CHH outside of ecdysozoans expands our understanding of the evolution and functional importance of the CHH-family. While the structural similarity is high, the sequences are quite distinct and even have a different number of disulfide bonds. There are other examples where the tertiary structures are more conserved than the amino acid sequence and, intriguingly, this is also the case for some signaling peptides. It is also notable that there are other examples of doppelganger toxins targeting this signaling system. Both venomous spiders, centipedes (Undheim et al. 2015), ticks and wasps (McCowan and Garb 2013) have convergently evolved toxins that mimic CHH peptides suggesting that the CHH and related peptides are functionally important in the prey.

Furthermore, using evolutionary trace analyses on the identified toxins and DREPs, we showed that the different precursor regions have district patterns of evolution: The toxin-containing regions are evolutionarily divergent compared to the signal and spacer regions in the toxin precursors, whereas the peptide-containing regions are more conserved in the DREPs. These contrasting patterns of evolution mirror the biological functions of the toxins and the DREPs. The toxins are adapting to receptors expressed in changing prey or predators, whereas the DREPs are maintaining activity in endogenous receptors that seldom change. We find that the combination of highly conserved signaling peptides, highly diverging toxins, and almost identical cone snail nerve ring paralogs make a very strong case for computationally assigning families of doppelganger toxins and their doppelganger-related peptides.

Additionally, our discovery of exon shuffling in Hairpin doppelganger toxins is, to our knowledge, the first confirmed example of a conotoxin that evolved by this mechanism. Conotoxins are grouped into superfamilies that share extensive signal sequence identity, and the toxins within these superfamilies have a common genetic architecture ranging from 1 to 6 exons (Phuong and Mahardika 2018). Here, we showed that cone snail Hairpin doppelganger toxins evolved by fusion of the first exons of contryphan genes with the endogenous cone snail DREP and thereby adopting the contryphan signal sequence. It is likely that exon shuffling has been a driving force in the evolution of multiple conotoxins but that this can no longer be detected. Several conotoxin superfamilies have distinct subfamilies that, while sharing a conserved signal sequence, contain different cysteine scaffolds and divergent precursor sequences. Exon shuffling could potentially explain the molecular evolution of this phenomenon.

Once recruited, DREPs can give rise to the evolution of toxin families with divergent pharmacologies that go beyond the structure and function of the original doppelganger toxins. For example, the C-superfamily of cone snail toxins evolved through recruitment of a somatostatin-related peptide, and most C-superfamily members are doppelganger toxins of this signaling peptide family (Koch et al. 2022; Ramiro et al. 2022). However, in some species these genes diversified to form the neurotensin receptor agonist, Contulakin-G, in *Conus geographus* (Craig et al. 1999)) and the nicotinic acetylcholine receptor antagonists αC-PrXA in *Conus parius* (Jimenez et al. 2007). The doppelganger toxins described here all structurally mimic the DREPs they evolved from, but future research may uncover more derived members of these doppelganger superfamilies.

We note that while we discovered several novel peptides in protostomes, we did not identify any novel signaling peptides in the zebrafish nor did we detect any orthologs of the five new DREP families outside of protostomes. However, our method was able to ‘rediscover’ several known zebrafish signaling peptides that share similarity with cone snail toxins, including oxytocin/vasopressin, insulins, and neuropeptide Y. While it is possible, that all doppelganger-related peptides have already been discovered in zebrafish, it is also possible that the evolutionary distance separating cone snails and zebrafish could cloud any homology and that other, more sensitive methods are needed to find any matching sequences. In this context, the phylogenetic distance between the venomous predator and its prey organisms is important to take into consideration.

In conclusion, this paper is a proof of concept for the systematic use of doppelganger toxins to discover unknown signaling systems. We anticipate that toxins from other organisms can be employed in a similar way using the generalizable approach described in this paper. Venomous and poisonous animals are not the only example of organisms that have evolved molecules to disrupt the behavior and physiology of another. Both parasites and pathogens are likely to use doppelganger toxins to manipulate their hosts to their advantage. Recently, several hormone-like sequences were detected in pathogenic viruses (Altindis et al. 2018; Huang et al. 2019). We propose that, in the future, the method described here can also be used to identify such yet-unknown genes in parasites and pathogens and their hosts.

## Materials and methods

### Phylogenetic analysis

COI, 12S, and 16S genes from diverse cone snails and *Californiconus californicus* were downloaded from NCBI. The three genes were individually aligned using MAFFT v7.487 and trimmed using trimAl v1.2 to remove all columns with gaps. The tree alignments were subsequently concatenated using FASconCAT-G v1.05. A maximum likelihood tree was constructed using IQ-TREE v 1.6.12 on a single threat. Based on the Bayesian information criterion the tree was constructed with TVM+F+I+G4 model of evolution. Bootstrap values were calculated with 1000 replicated using IQ-TREE’s UFboot method.

### Transcriptome sequencing

Specimens of *A. californica* were ordered from the National Resource for Aplysia at the University of Miami, FL, USA. Animals were anesthetized as previously described (Zhao et al. 2009). The following ganglia and non-neuronal tissues were dissected from two specimens: abdominal ganglion, cerebral ganglion, right and left pedal ganglia, right and left pleural ganglia, right and left buccal ganglia, nerves (connecting the abdominal with the pedal ganglia), spermatheca, opaline gland, purple gland, salivary gland, and foot clip. The venom gland of a single specimen of *Conus furvus* was also dissected for sequencing. All other venom gland datasets were retrieved from public repositories as described below. Tissues were stored in RNAlater (ThermoFisher Scientific) at −80°C until further processing. Total RNA was extracted using the Direct-zol RNA extraction kit (Zymo Research), with on-column DNase treatment and an additional wash step after the first purification, according to the manufacturer’s instructions. Library preparation and sequencing were performed by the University of Utah High Throughput Genomics Core Facility as previously described for different cone snail tissues (Koch et al. 2022).

### Prey databases preparation

The prey databases were built through a generalizable pipeline with different inputs for the *Danio rerio, Aplysia californica*, and annelid (*Platynereis dumerlii* and *Capitella teleta*) databases. These species were chosen because of the availability of genomic and transcriptomic material, phylogenetic relationship to cone snail prey, and status as model organisms.

The *Aplysia* database was formed by downloading all proteins of *Aplysia californica* from the NCBI Protein database with the query ““Aplysia californica”[porgn: __txid6500]” in December 2021 (27,891 sequences). The redundant sequences were removed using cd-hit at a similarity level of 95 % and proteins with signal sequences were extracted using SignalP 6.0 (2,649 sequences). To account for novel proteins that were not predicted from the genome, we added secreted proteins from several assembled transcriptomes from *A. californica* (GBCZ00000000.1, GBDA00000000.1, GBAV00000000.1, GBBV00000000.1, GBBG00000000.1, GBBE00000000.1, GBBW00000000.1, GBAQ00000000.1, GAZL00000000.1). Potential open reading frames of at least 50 amino acids were extracted with getorf and clustered at a similarity level of 90 % using cd-hit. All potential methionine start-sites were assessed with SignalP6.0 and sequences with signal peptides were kept. In case several start-sites from a single transcript were predicted to have a signal sequence, only the longer sequence was kept (8,997 sequences). The secreted sequences from the NCBI protein database and the transcriptomes were concatenated and clustered with cd-hit at 99 % similarity (10,039 sequences). Proteins with transmembrane domains were removed with TMHMM2.0. Sometimes signal peptides are erroneously predicted as signal sequences in TMHMM, and thus sequences predicted to have a single transmembrane domain in the first 30 amino acids were kept for subsequent analyses (8,515 sequences). We noticed that many of the secreted proteins were already annotated as enzymes in UniProt. To remove these enzymes, we downloaded all. catalytic proteins from UniProtKB from mollusks with the following terms taxonomy: ““Mollusca (9MOLL) [6447]” AND goa:(“catalytic activity [0003824]”)”. Enzymes were removed from the database with mmseqs at an e-value of 1E-10 resulting in 7,009 secreted sequences that, in principle, include all secreted neuropeptides and peptide hormones.

For the zebrafish database, we downloaded all proteins from the NCBI Protein database with the query ““Danio rerio”[porgn:__txid7955]” and the altorfs from https://www.roucoulab.com/en/downloads.html based on the Ensembl zebrafish annotation Zv9.97 (total 177,106 sequences). Using the approach above we identified 5,929 secreted proteins. These were supplemented with 11,562 secreted sequences identified from three assembled transcriptomes (GDHQ0000000.1, GDQQ0000000.1, and GFIL00000000.1). Following removal of sequences with transmembrane domains (11,374 seqs) and similarity to chordate enzymes (Uniprot search terms “taxonomy:”Chordata (9CHOR) [7711]” goa:(“catalytic activity [0003824]”)”) the final zebrafish database consisted of 9,328 sequences.

The final prey database was built from 32,117 sequences downloaded from the NCBI Protein database with the search term: ““Capitella teleta”[porgn:__txid283909]”, of which 2,483 sequences had a predicted signal sequence. We also added 11,729 secreted protein sequences identified from four transcriptomes of the annelid *Platynereis dumerlii* (GBZT00000000.1, HALR00000000.1, HAMN00000000.1, and HAMO00000000.1), as there were no assembled transcriptomes available at the time of the search. Following the removal of transmembrane proteins (12,127 sequences) and enzymes (Uniprot search terms: “taxonomy: “Annelida [6340]” AND goa:(“catalytic activity [3824]”)”) the final annelid database consisted of 10,659 secreted proteins.

Accession numbers of all SRA datasets used in this paper can be found in **Supplementary file 12**. Code is available from https://github.com/Thomaslundkoch/toxmims.

### Venom database preparation

We downloaded 92 transcriptomes from 45 different species of cone snails representing diverse clades with different prey preferences from NCBI (SRA accession numbers listed in **Supplementary file 12**. These were assembled as previously described (Koch et al. 2022). The assembled venom gland transcriptomes were processed individually in a process identical to the transcriptome of *A. californica* with slightly different settings (code is available from https://github.com/Thomaslundkoch/toxmims). The open reading frames were only clustered with cd-hit at 100 % identity and pooled in the end.

We noticed that SRR1544627 and SRR11807494 appeared contaminated by the snails’ nervous tissue (based on an unusually high number of different neuropeptides in the transcriptomes). Consequently, these were not included in the subsequent analyses. Contamination of the venom gland by nervous tissue typically happens if the venom gland is not properly separated from the area of the pharynx that is closely connected to the circumesophagal nerve ring (Safavi-Hemami et al. 2016).

### Venom homology search

The proteins from the ‘prey’ databases were used to query the combined venom database from cone snail venom gland transcriptomes with BLASTP. We used a word size 2 and e-value 1e-2 in the searches. A total of 515 sequences from the *Aplysia* database had significant hits, 675 in the zebrafish database, and 1020 in the Annelid database. For each hit in the prey databases, we created a multiple sequence alignment with the venom blast hits with TPM above 10. The alignments were then visually inspected. Alignments that showed a characteristic doppelganger toxin pattern (a combination of highly conserved and diverse amino acid residues in a potential mature peptide region) and putative processing sites were further analyzed by searching for orthologs in closely related species using BLASTP against the NCBI non-redundant protein database and TBLASTN against TSAs of other members of the prey phylum.

### Evolutionary rate analysis

Sequences were aligned using MAFF v7.487 and the evolutionary rates were calculated using rate4site. The evolutionary rates were plotted using a sliding window of 5 amino acids. The boxplots were built from the evolutionary rates of the peptide and pro-peptide regions as shown in the alignment figures (the likely processing sites were left out of the analysis) and compared using Wilcoxon rank-sum test. Rate4site scores have been shown to be strongly correlated with and directly comparable to dN/dS values (Sydykova and Wilke 2017).

### Gene structure analysis

We identified the location, size, and phases of introns using the online version of Splign. The mRNA was obtained from the respective transcriptomes, and the corresponding genomic segment was identified using tblastn with standard setting.

### Structural prediction and comparison

We obtained structural predictions of the toxins and putative signaling peptides using a combination of AlphaFold2 neural network and MMSeqs2 to obtain a multiple sequence alignment. These are combined in ColabFold, where the full precursor sequences were used as the query sequence. To avoid similarities due to a common template, we performed a structural prediction without templates. The best of five Amber relaxed models was selected. Following prediction, we removed the signal sequence, pro-peptide regions, and processing sites to obtain a model of the mature toxin/ signaling peptide. As a rule of thumb pLDDT values above 90 are expected to be modeled with high accuracy; 70-90 are modeled well with a generally good backbone prediction. pLDDT from 50-70 should be handled with caution. Values below 50 are often represented as a helix, this should, however, not be interpreted, and it is a reasonably strong indicator of disorder (Jumper et al. 2021).

We used the protein structural comparison server DALI to compare the predicted toxin and signaling peptide structures to all available protein structures in PDB and different species subsets of the AlphaFold database.

### Clustering analysis

Clustering analysis was performed using CLANS (Frickey and Lupas 2004), which randomly initializes the individual sequences as nodes and performs an all-against-all BLASTP. The negative logarithm of the BLAST p-values is transformed into an attractive force in addition to a uniform repulsive force between the nodes. We used the BLOSUM62 scoring matrix using the web tool https://toolkit.tuebingen.mpg.de/tools/clans. The clustering was initially done in 3D and collapsed to 2D for >300,000 rounds, at which point the clustering had converged.

## Acknowledgments

We would like to thank the National Resource for Aplysia at the University of Miami, USA, the High Throughput Genomics Core Facility at the University of Utah, USA for library preparation and transcriptome sequencing, and Maren Watkins for assistance with retrieving phylogenetic marker genes.

